# A 3D Image Segmentation Study of U-Net on CT Images of the Human Aorta with Morphologically Diverse Anatomy

**DOI:** 10.1101/2024.10.02.616348

**Authors:** Jiasong Chen, Linchen Qian, Pengce Wang, Christina Sun, Tongran Qin, Asanish Kalyanasundaram, Mohammad Zafar, John Elefteriades, Wei Sun, Liang Liang

**Affiliations:** Department of Computer Science, University of Miami, FL; Sutra Medical Inc, CA; Yale University Medical School, CT

## Abstract

For the machine learning -assisted diagnosis of cardiac diseases, such as thoracic aortic aneurysm, the geometries of the heart and blood vessels need to be reconstructed from medical images, which is usually done by image segmentation followed by meshing. In this study, we applied U-Net (2D and 3D versions), a deep neural network with a U-shaped architecture, to segment human aorta from CT images. From our experiments, we have the following observations: (1) 2D U-Net, which segments each of the 2D slices of a 3D CT image independently, produced erroneous fragments (e.g., missing part of the aorta) and boundaries (i.e., aortic walls) in 3D; (2) 3D U-Net, which does segmentation in 3D regions of a 3D CT image, performed much better than 2D U-Net. We also observe the major weakness of the 3D U-Net: the reconstructed geometries of the aortic wall had large errors (measured by HD95) for some cases. The 3D U-Net in this study serves as a baseline for developing more advanced architectures of deep neural networks for more accurate geometry reconstruction of human aorta.

## 1. INTRODUCTION

Cardiovascular diseases (CVDs) are the leading cause of death globally. In the United States, CVDs, including heart disease and stroke, are the primary causes of mortality [1,2]. Cardiac imaging plays a crucial role in diagnosing CVDs. Various medical imaging techniques, including X-ray, computed tomography (CT), magnetic resonance imaging (MRI), echocardiography (Echo), positron emission tomography (PET), and single-photon emission computed tomography (SPECT) [3], are employed to visualize and assess the structure and function of the heart and blood vessels.

Deep learning, i.e., machine learning with deep neural networks, has proven to be an essential tool to automate medical image analysis tasks, including image segmentation [4–8], image classification [9–13], image registration [14,15], image enhancement [16,17], and image generation [18–20]. Cardiac image segmentation techniques can provide detailed structural information about the heart and blood vessels, while registration techniques perform local functional analysis, aiding in both diagnosis and treatment planning [21]. Chen et al. [4] conducted a systematic review of deep learning methodologies across three prevalent imaging modalities (Ultrasound, CT, and MRI), focusing on the assessment of various cardiac anatomical structures. Their study shows that contemporary deep learning techniques enhance segmentation performance by utilizing advanced building blocks for feature extraction, designing sophisticated loss functions, implementing multi-task learning, and applying regularization through external constraints.

The aorta is the body’s largest artery, responsible for distributing oxygenated blood throughout the systemic circulation [22]. Aortic aneurysm disease ranks consistently in the top 20 causes of death in the U.S. population [23]. Thoracic aortic aneurysm (TAA) is a bulging of the thoracic aortic wall and it is a leading cause of death in adults [24]. TAA prevalence is relatively high, estimated to be around 1% of the general population [25,26]. TAA dissection happens when a tear forms in the inner layer of the aorta, leading to the separation of the aortic wall layers [27]. TAA rupture happens when the aortic wall tissues in all the layers break apart under high blood pressure. The rupture or dissection of TAA will lead to death if left untreated. For high-risk aneurysm patients, elective repair will be recommended to replace the aneurysm (i.e., part of the native aorta) with a graft in order to prevent the aneurysm from rupturing or dissecting. Currently, 3D CT imaging is the gold standard for diagnosing TAA. The risk of rupture and dissection is partially determined by the aorta geometry [28] that can be obtained from 3D CT images.

To obtain the aorta geometry of a patient from 3D CT images, automated image segmentation methods have been developed, which are mostly based on a specific architecture of the deep neural network, known as the U-Net [29]. For example, Cao [5] developed three convolutional neural network models based on the U-Net to simultaneously segment the entire aorta, true lumen (TL), and false lumen (FL), demonstrating the high accuracy and efficiency of U-Net-based models for CT image segmentation of the aorta with type B aortic dissection. Cheung et al. [6] employs a 2D U-Net to segment the coronary arteries with high accuracy. Saitta [30] developed a fully automated pipeline that employs a 3D U-Net for thoracic aorta segmentation, subsequently integrating the segmentation results into a computational geometry processing pipeline that generates relevant aortic metrics for TEVAR planning.

The performance of image segmentation is also affected by non-architectural factors, including preprocessing, training, and inference techniques. Currently, nnU-Net [31] achieves competitive segmentation performance across various datasets by implementing an automatic training framework based on the vanilla U-Net architecture. nnU-Net can automatically adapt the simple U-Net architecture to the specific image dataset and available hardware resources.

In this study, the objective is to train deep neural network models to accurately and effectively segment the whole aorta (including the aortic root) with anatomically diverse structures from 3D cardiac CT images. For this segmentation task, we explored two versions of the nnU-Net: (1) 2D U-Net, which segments each of the 2D slices of a 3D CT image independently; (2) 3D U-Net, which divides a 3D image (e.g., 512×512×512) into smaller 3D regions (e.g., 128×128×128) and does segmentation in each of the 3D regions. By dividing a 3D image into smaller regions, the 3D U-Net can fit into the VARM of a single GPU during training.

## 2. MATERIALS AND METHODS

The main objective of this study is to evaluate the segmentation performance of nnU-Net on datasets that encompass a diverse range of aortic shapes and conditions.

### 2.1. Datasets

We combine two de-identified datasets for this study, which consists of a total of 74 patients. The first dataset is from Yale University – New Haven Hospital, which has 47 patients. The CT images in this Yale dataset include images of patients without TAA and those who have TAA and Type A aortic dissections at multiple imaging dates. The annotations for the dataset were carried out by three experienced experts, with 1 to 1.5 hours to annotate an image. The annotations contain two substructures: the aorta (AO) and the aortic root. The second dataset is derived from a publicly available dataset, known as HOCM [32]. It comprises 27 3D CT images obtained using a Siemens SOMATOM Definition Flash scanner. All 27 patients in this dataset had undergone septal myectomy surgery. In the combined dataset of 74 cases, the median voxel spacing is 1.00×0.88×0.88 mm, and the median image size is 512×512×462. Additional information of the datasets is provided in Table 1.

**Table 1.** Some details of the CT datasets.

| Dataset | Number | Age | Image Size | Median Voxel Spacing | Substructures in Annotations |
| --- | --- | --- | --- | --- | --- |
| HOCM | 27 | 38-76 | 512×512× (275-571) | 0.25×0.25×0.5mm | AV, MV, AO, LA, LV, Myocardium, and excised myocardium |
| Yale | 47 | >30 | 512×512× (62-2092) | 0.742×0.742×0.625mm | AO, Aortic root |

Examples of the images and annotations (i.e., reconstructed geometries of the aorta) are illustrated in Figure 1 and Figure 2 using the 3D slicer software, which provides a comprehensive visualization of the cardiac structures for annotation. It can be seen that aortic aneurysm w/o dissection can lead to distinct shape changes, compared to normal aorta shapes.

**Figure 1.**
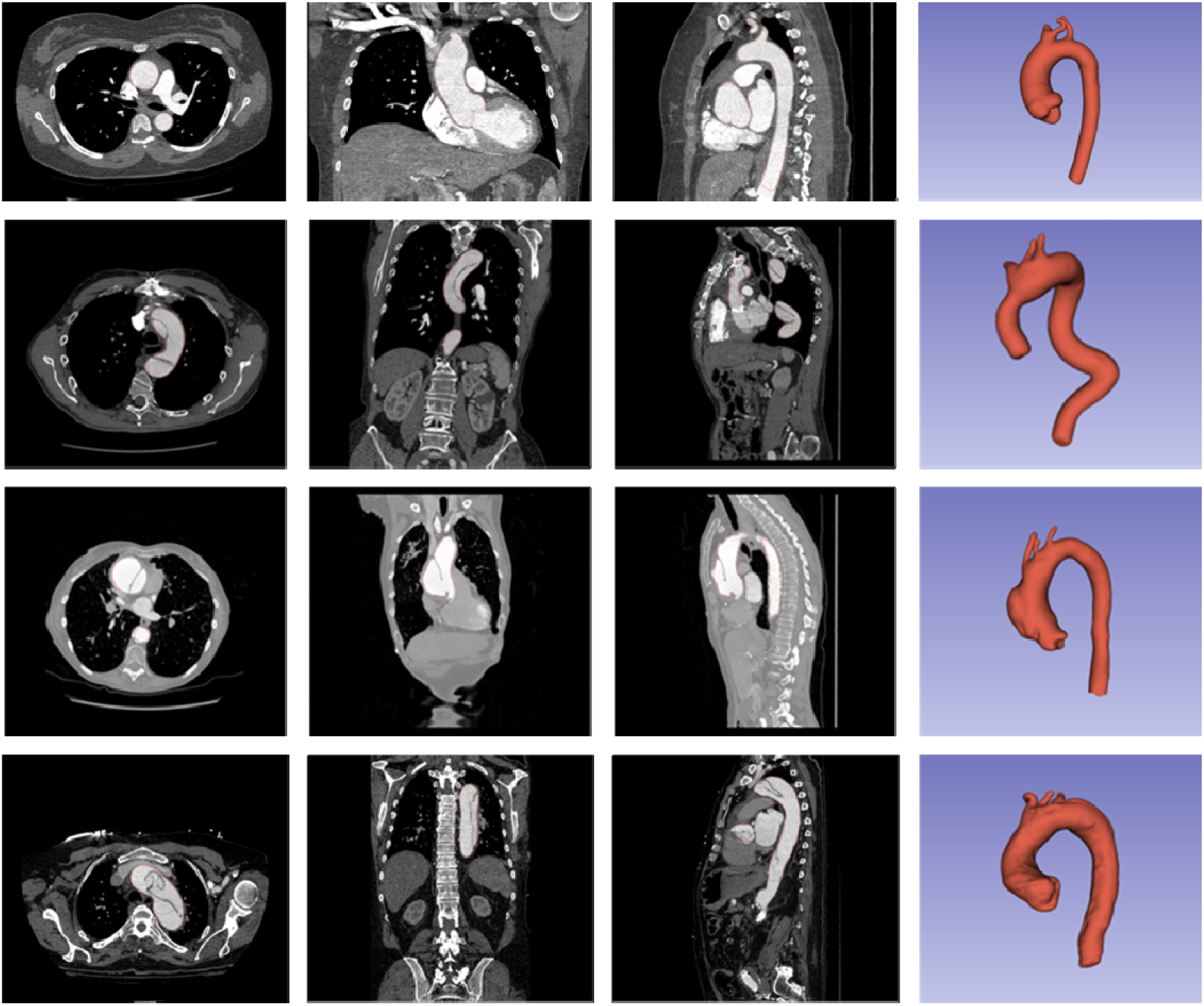
The cardiac CT images of four human subjects are presented in multiple views (axial, sagittal, and coronal), and the 3D annotations are shown in the rightmost column. The first row illustrates the image and normal shape of the aorta. The last three rows show examples of aortic shapes affected by aneurysms w/o dissections.

**Figure 2.**
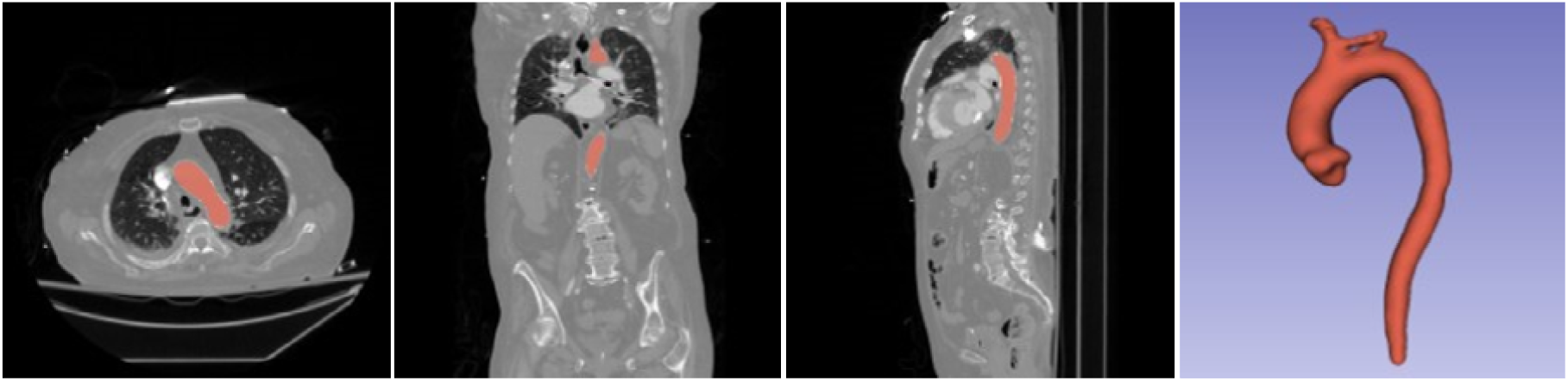
A zoom-in view of the image and aorta shape of a human subject without aortic aneurysm. aneurysm w/o dissection can lead to distinct shape changes, compared to normal aorta shapes.

### 2.2. Network Structure

The nnU-Net framework was chosen to carry out the segmentation task. nnU-Net employs a simple U-Net architecture with a well-optimized pipeline to achieve state-of-the-art segmentation performance. Its network structure includes an encoder path for down-sampling and a decoder path for up-sampling. The network has a U-shaped configuration, connecting the encoder and decoder paths through skip connections, which enhances segmentation performances. Various U-Net configurations can be integrated within the nnU-Net architecture. The original architecture is composed of multiple resolution stages, with each stage in both the encoder and decoder comprising two computational blocks. Each computational block follows the sequence of convolution, instance normalization, and Leaky ReLU activation.

In this study, both 2D U-Net and 3D U-Net were selected for integration into the nnU-Net framework. Figure 3 shows the details of 2D U-Net and Figure 4 shows the structure of 3D U-Net. For the 2D U-Net, a 3D image of the aorta images is divided into 2D slices, each of the 2D slices is an input to the U-Net model. This approach results in a loss of spatial information in the slicing direction. Utilizing a 3D U-Net for the analysis of 3D aorta data is a judicious approach; however, conventional 3D segmentation techniques may exhibit suboptimal performance in the presence of anisotropic datasets [31,33]. Furthermore, owing to limitations in GPU memory, nnU-Net employs patch-based training for datasets containing large images, i.e., dividing a 3D image into smaller 3D regions, and each of the 3D regions is used as an input to the 3D U-Net model. This 3D patch-based approach may hinder the 3D U-Net’s ability to capture sufficient contextual information necessary for effective segmentation.

**Figure 3.**
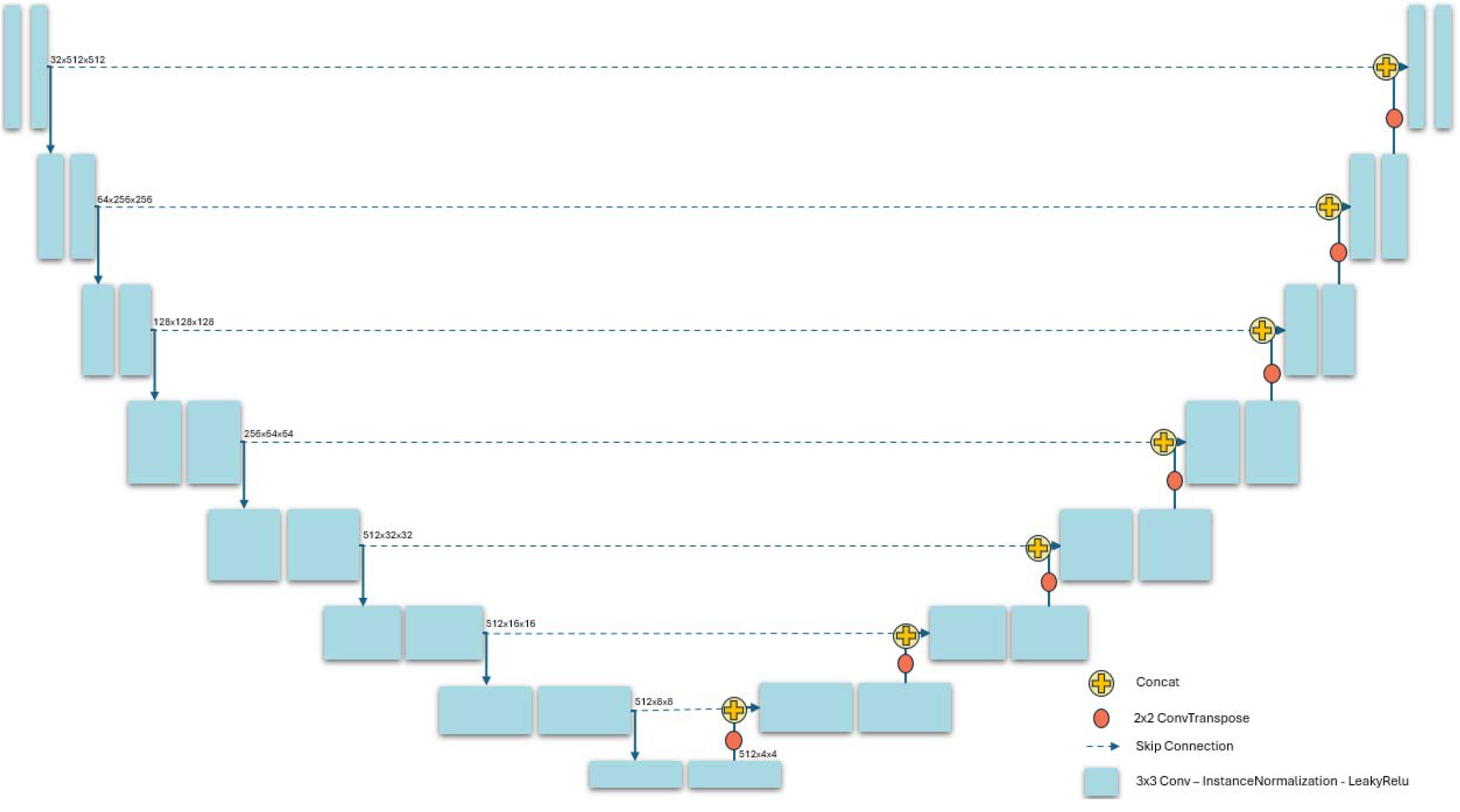
Network structure of 2D U-Net

**Figure 4.**
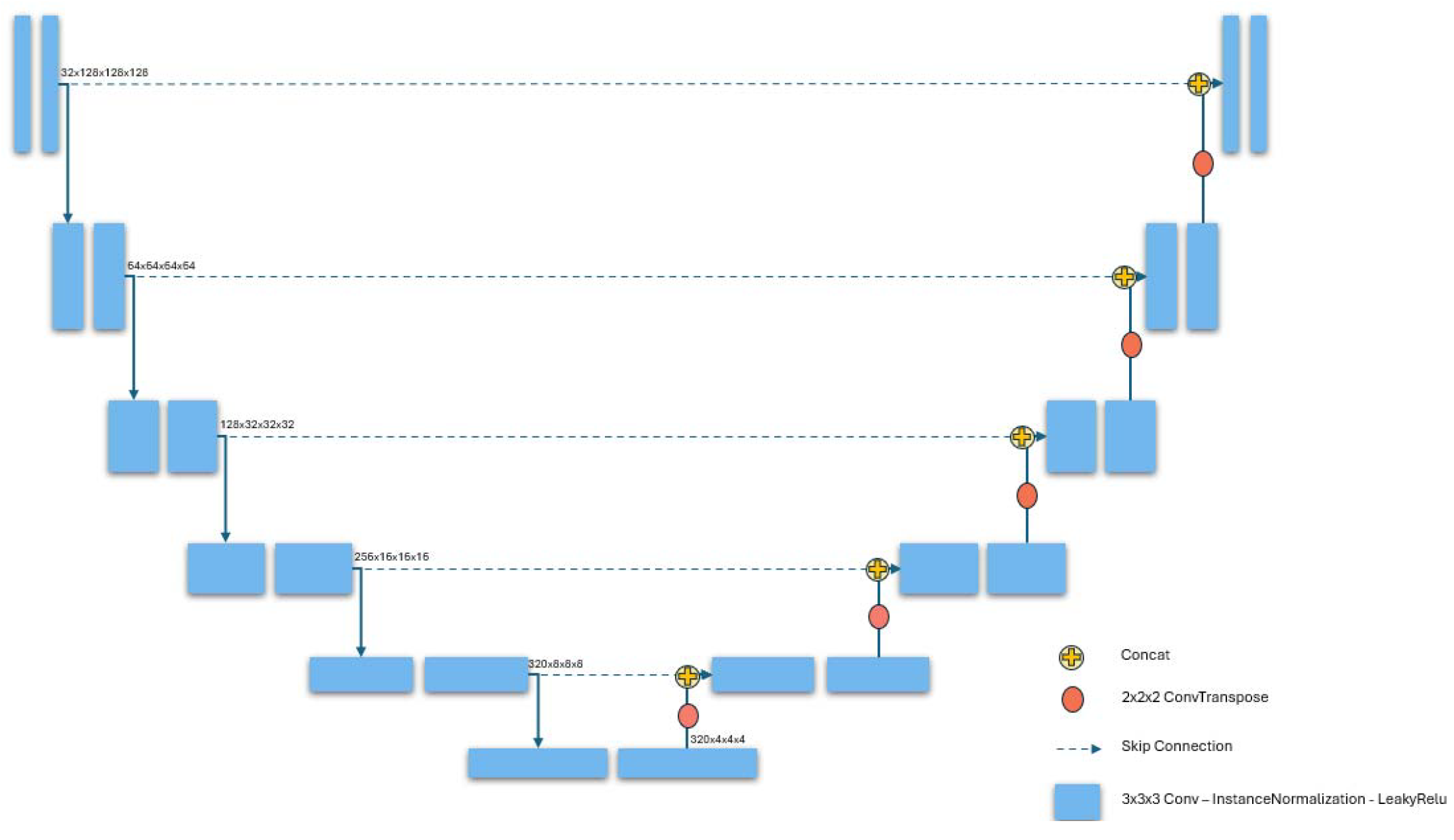
Network structure of 3D U-Net

## 3. EXPERIMENTS AND RESULTS

### 3.1. Annotation Refinement

For this study, we focused solely on the segmentation of the aorta and aortic root. The data annotations are further processed before being used by the image segmentation models, which has several key steps, including segmentation refinement, removing small arterial branches, and standardizing the arterial branches extending from the aortic arch. These processing steps ensured more accurate and consistent annotations of the aorta for further analysis. Some of the data from the HOCM dataset lack annotations of the descending aorta. Three experienced experts conducted and validated the annotation refinement specifically for these cases. The annotation of the aorta terminates at the level of the fourth lumbar vertebra, where it bifurcates into the right and left common iliac arteries. In the annotation process, the left and right coronary arteries were excluded. Additionally, three arterial branches from the aortic arch—the brachiocephalic artery, the left common carotid artery, and the left subclavian artery—were trimmed to a uniform length of approximately 25mm. Small arterial branches on the descending aorta were also removed. All modifications to the segmentation were verified and reached a consensus among the experienced experts. An example is shown in Figure 5.

**Figure 5.**
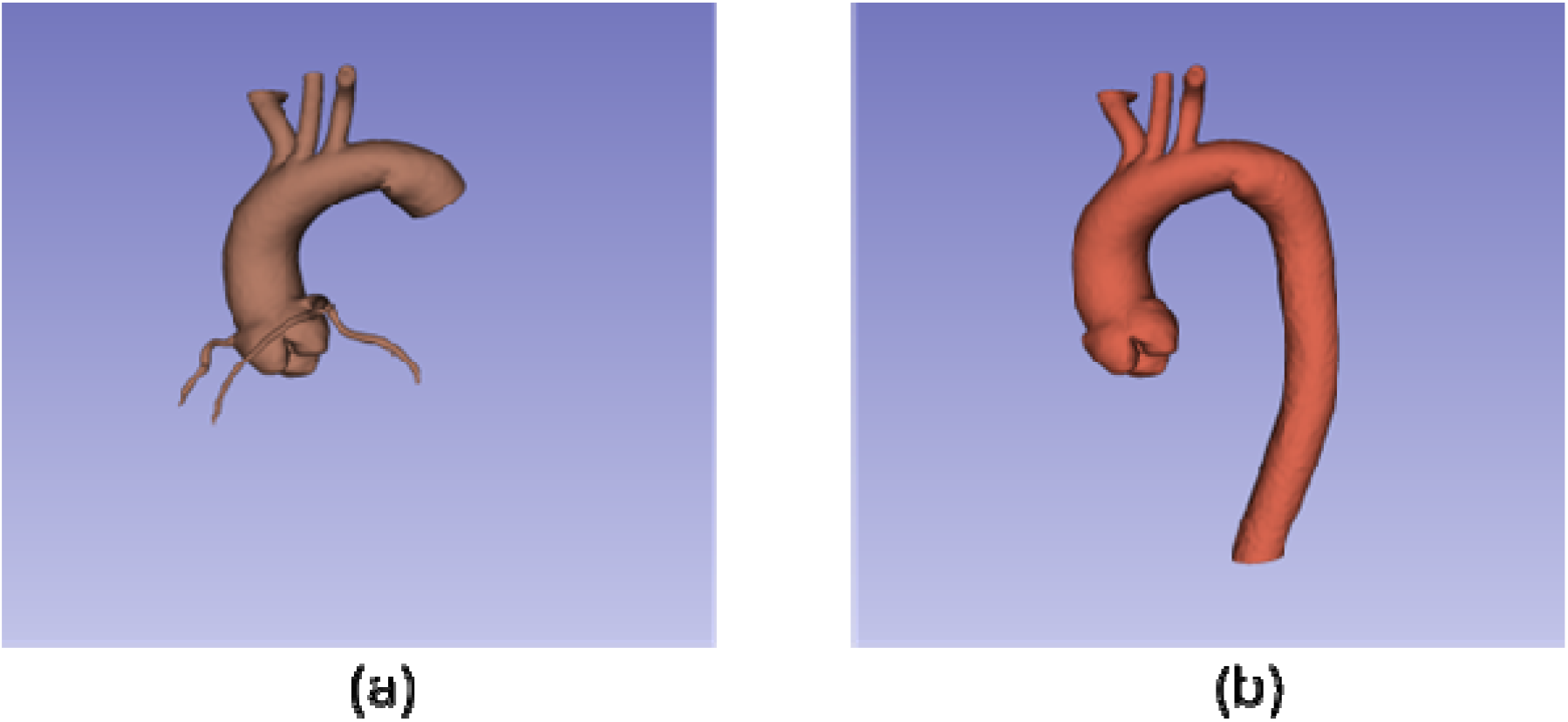
An example of annotation refinement: (a) Before refinement, (b) After refinement.

### 3.2. Data Processing

In the nnU-Net training pipeline, several data processing operations are performed on the CT images, including image cropping, resampling, and data normalization. First, the CT images are cropped to remove pure black background. Next, the CT images are resampled using a median voxel spacing of 0.88×0.88×1.00 mm with third-order spline interpolation. After resampling, each CT image is normalized by clipping the intensity values to fall within the range of the 0.5% to 99.5% percentiles of all intensity values in the dataset. This is followed by z-score normalization based on the global mean and global standard deviation to ensure consistent intensity scaling across the dataset.

### 3.3. Experiment Setup

The dataset, consisting of 74 cases, was randomly divided into 68 training samples and 6 testing samples. To enhance the model’s generalization capabilities, five-fold cross-validation was employed during the training process. This allows for more robust evaluation and optimization of nnU-Nets. During the cross-validation process, the 68 training samples were further randomly split into 54 for training and 14 for validation purposes. The best model configuration identified from this process was then applied for model evaluation on the 6 testing samples. Throughout the training process, various data augmentation techniques were applied to enhance model robustness, including rotation, scaling, Gaussian noise, Gaussian blur, brightness processing, and contrast adjustment. These augmentations aimed to improve the model’s ability to generalize across different conditions and variations in the image data.

Both the 2D U-Net (Figure 3) and the 3D U-Net (Figure 4) were selected for integration within the nnU-Net framework. For the training of the 3D U-Net, the original CT images were divided into 3D patches of size 128×128×128 to accommodate computational requirements. A batch size of 20 was used during training. The network consists of a total of 6 stages with different resolutions. In each stage, there are two computational blocks for both the down-sampling and up-sampling paths. Each convolutional block includes a convolutional kernel of size 3 × 3 × 3, followed by an instance normalization (IN) layer and a Leaky ReLU activation. The down-sampling process ultimately produces 320 feature maps with a size of 4×4×4. Subsequently, the feature maps are up-sampled through different stages until the feature map size is restored to the size of the input patch.

The 2D U-Net utilized a patch size of 512×512, with a batch size set to 12. Each 3D CTA image was sliced into individual 2D slices, which were then fed into the 2D U-Net. The model comprises 8 stages with different resolutions. The down-sampling process culminates in 512 feature maps, each with a size of 4×4. Subsequently, these feature maps are up-sampled through different stages until the feature map size is restored to the size of 512×512. This allows the network to effectively learn and segment the aorta and aortic root region from the 2D slices.

The training loss ℒ for the model is a combination of the Dice coefficient similarity (DSC) loss ℒ_*Dice*_ and the standard cross-entropy loss ℒ_*CE*_, as shown in Eq. (1) and Eq. (2). A total of 1000 epochs were conducted during training. For optimization, stochastic gradient descent with an initial learning rate of 0.01 and Nesterov momentum set at 0.99 was employed. The learning rate was gradually reduced to zero using a “polyLR” schedule throughout the training process [34,35]. The experiments were conducted on a server with Nvidia H100 GPUs, each with 80 GB of memory, ensuring efficient processing and training of the models.

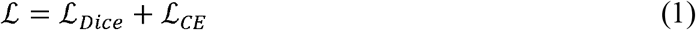

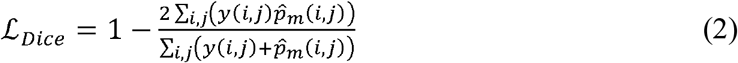

### 3.4. Experiment Results

To evaluate the segmentation performance of nnU-Net (2D and 3D), we employed several metrics: the Dice Score Coefficient (DSC), the 95th percentile of the Hausdorff Distance, Jaccard Index, precision, and accuracy. These metrics provide a comprehensive assessment of a model’s ability to capture and segment the aorta and aortic root region. The <u>Dice Score Coefficient</u> assesses the overlap between the predicted segmentation and the ground truth, providing a measure of their similarity. The <u>Hausdorff Distance</u> quantifies the maximum distance between the boundaries of the predicted segmentation masks and the ground truth, indicating how far apart the two boundaries are. Hausdorff Distance at the 95th percentile (HD95) is usually used to ignore outliers. Additionally, the <u>Jaccard Index</u> evaluates the similarity of the two segmentation masks by comparing their intersection and union, offering another perspective on the accuracy of the segmentation results. The <u>precision</u> metric measures the proportion of pixels predicted as part of the aorta by the model that are indeed true aorta pixels in the ground truth segmentation. Conversely, the <u>recall</u> metric quantifies the amount of actual positive pixels in the ground truth segmentation masks that were correctly identified as positive by the segmentation model. The <u>accuracy</u> metric calculates the proportion of correctly classified pixels in the predicted segmentation masks relative to the total number of pixels in the segmentation masks.

The quantitative evaluation of the performance is shown in Table 2, using the abovementioned metrics. This comparison allows for an assessment of how each version of the nnU-Net performs in accurately segmenting the aorta and aortic root region. For each evaluation metric, the table displays the minimum, maximum, and average values. Generally, larger values indicate better segmentation performance for most metrics assessed, except the HD95 where smaller values indicate better segmentation performance. nnU-Net 3D outperforms nnU-Net 2D on average for DSC, HD95, Jaccard Index, Precision, and Accuracy. nnU-Net 2D has better average recall.

**Table 2.**
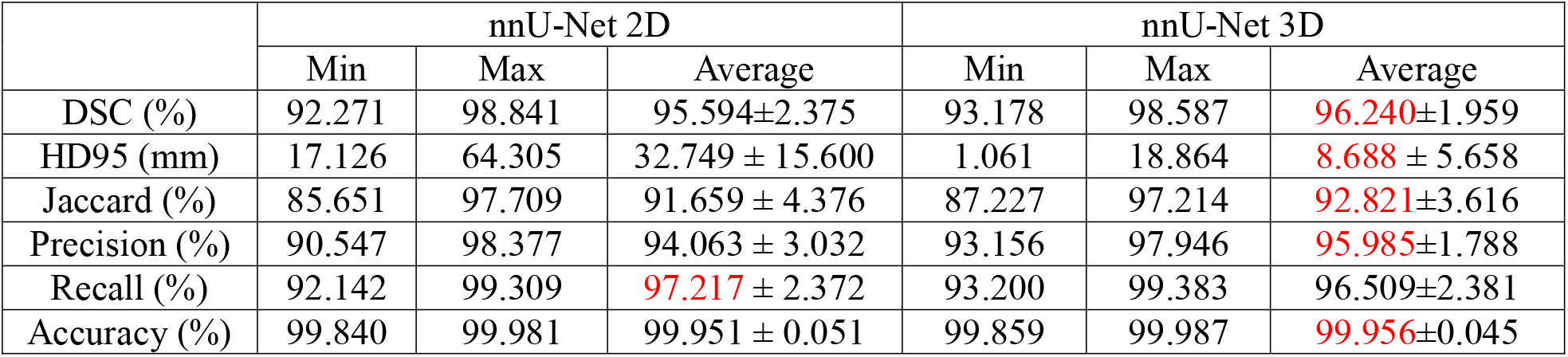
Performance of nnU-Net with 2D U-Net and 3D U-Net.

|  | nnU-Net 2D |  |  | nnU-Net 3D |  |  |
| --- | --- | --- | --- | --- | --- | --- |
|  | Min | Max | Average | Min | Max | Average |
| DSC (%) | 92.271 | 98.841 | 95.594±2.375 | 93.178 | 98.587 | 96.240±1.959 |
| HD95 (mm) | 17.126 | 64.305 | 32.749 ± 15.600 | 1.061 | 18.864 | 8.688 ± 5.658 |
| Jaccard (%) | 85.651 | 97.709 | 91.659 ± 4.376 | 87.227 | 97.214 | 92.821±3.616 |
| Precision (%) | 90.547 | 98.377 | 94.063 ± 3.032 | 93.156 | 97.946 | 95.985±1.788 |
| Recall (%) | 92.142 | 99.309 | 97.217 ± 2.372 | 93.200 | 99.383 | 96.509±2.381 |
| Accuracy (%) | 99.840 | 99.981 | 99.951 ± 0.051 | 99.859 | 99.987 | 99.956±0.045 |

Figure 6 shows examples of the segmentations using the two versions of the nnU-Net. Although the segmentations from nnU-Net 2D and nnU-Net 3D have similar overall shapes, there are noticeable differences.

**Figure 6.**
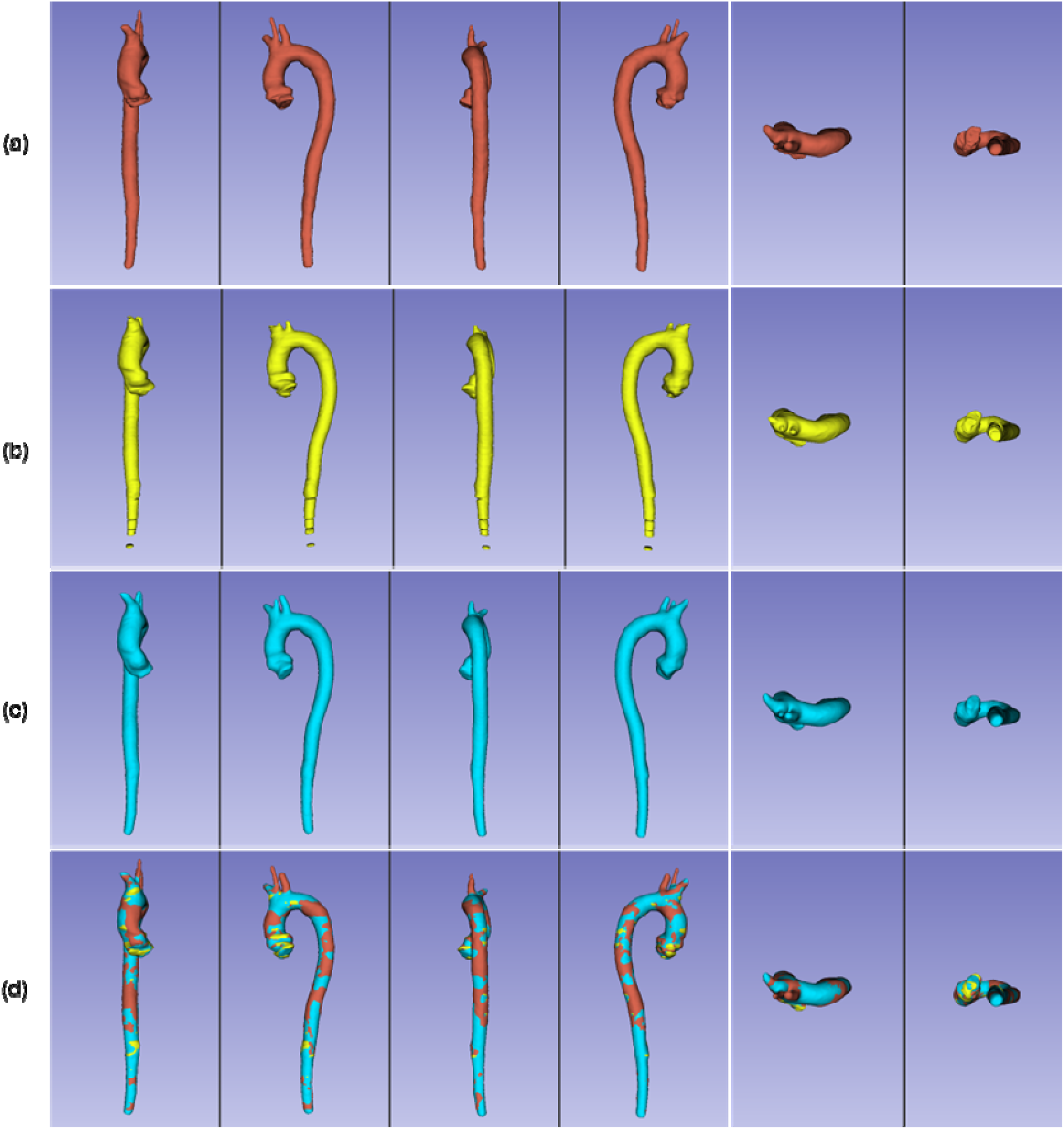
Segmentation examples. The row-(a) shows the ground-truth segmentation of the aorta in a patient. The row-(b) shows the segmentation result of the 2D nnU-Net. The row-(c) shows the segmentation result of the nnU-Net3D. The row-(d) show all of the segmentations.

Figures 7 and 8 provide a detailed comparison of nnU-Net 2D and nnU-Net 3D from multiple views, including axial, sagittal, coronal, and 3D perspectives. The first row displays the axial view segmentations of arteries on the aortic arch. The nnU-Net 3D demonstrates superior segmentation performance compared to the nnU-Net 2D. The 3D kernel is better equipped to capture volumetric information from the scans, allowing for more accurate segmentation of the predefined lengths of arteries on the aortic arch, which aligns closely with the ground truth segmentation. The second and third rows present the axial view segmentation results for the upper and middle sections of the aorta. The upper aorta axial view encompasses both the ascending and descending aorta region, while the middle aorta axial view only contains the descending aorta region. Both nnU-Net 2D and nnU-Net 3D perform well in these sections, effectively capturing the anatomical features and accurately segmenting the aorta area. The fourth row reveals that the nnU-Net 2D fails to identify the bottom section of the aorta, while the nnU-Net 3D successfully detects the aortic area. The last three rows display segmentation results in coronal and sagittal views. Both nnU-Net 2D and nnU-Net 3D perform well in aorta segmentation, but the nnU-Net 3D yields smoother edge contours that align more closely with the ground truth segmentation mask, indicating better overall performance in capturing the shape of the aorta and aortic root.

**Figure 7.**
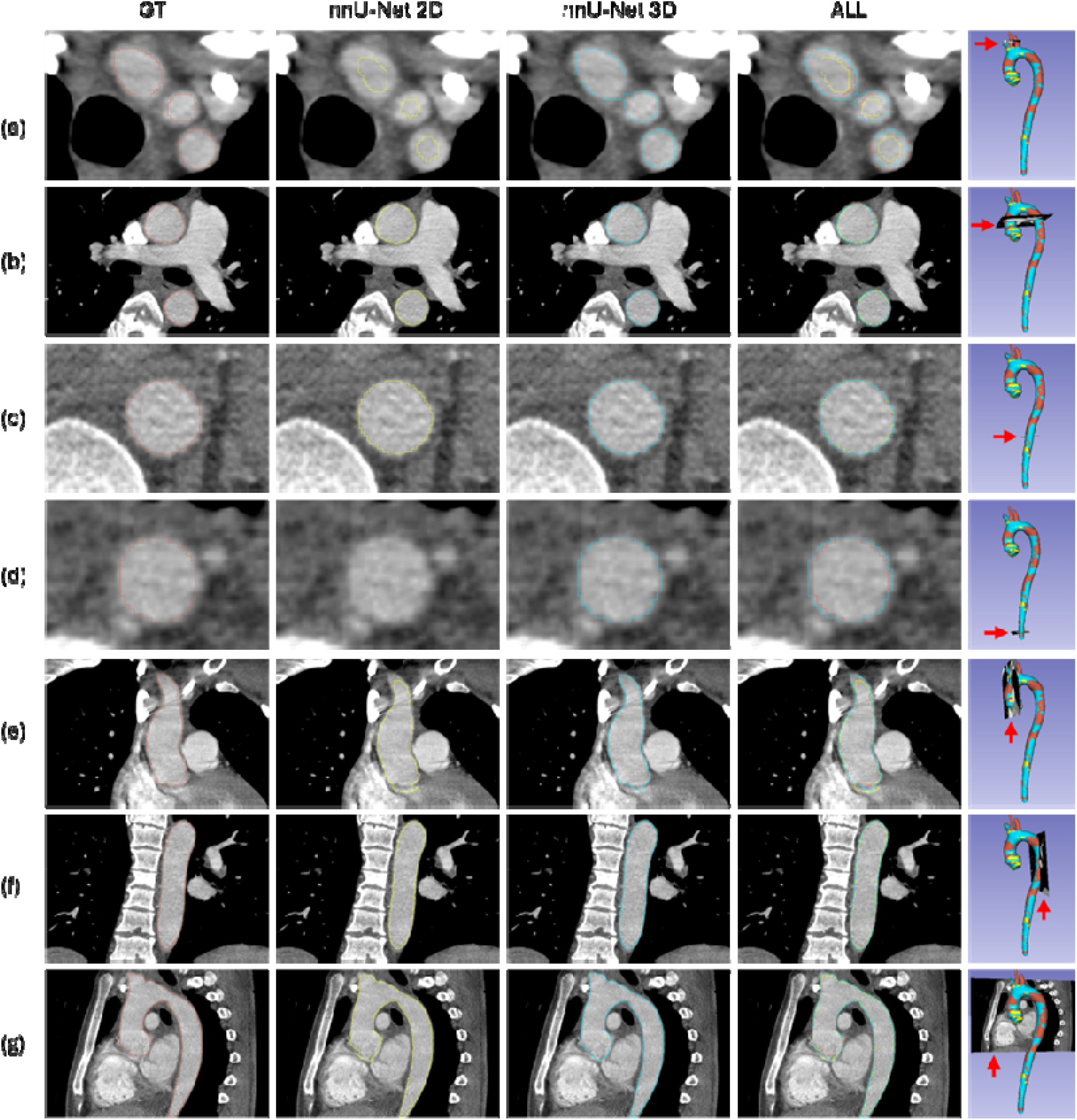
Comparison of nnU-Net 2D (Yellow), nnU-Net 3D segmentation (Blue), and ground-truth (GT) segmentation (Red).

**Figure 8.**
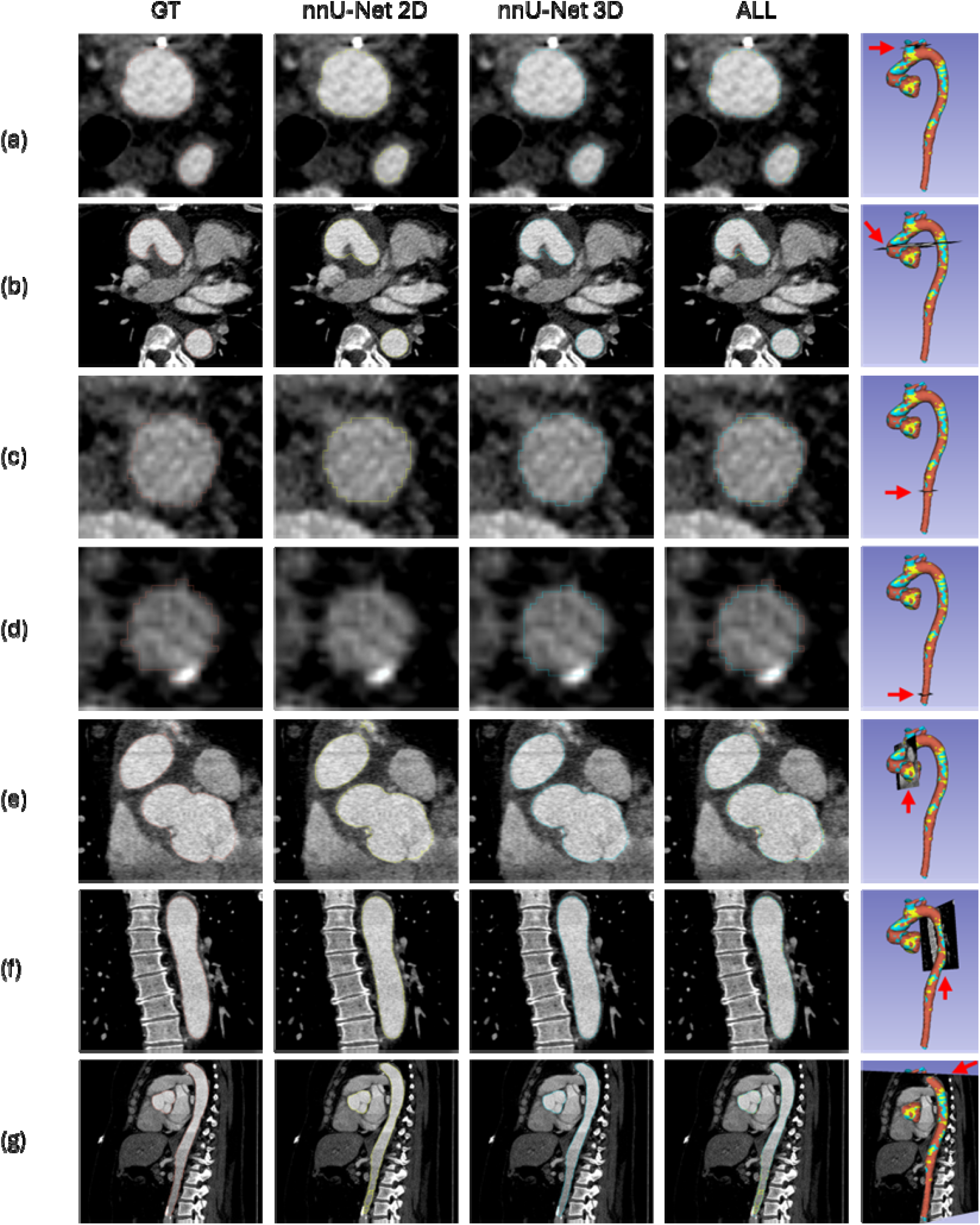
Comparison of nnU-Net 2D (Yellow), nnU-Net 3D segmentation (Blue), and ground-truth (GT) segmentation (Red).

By zooming into Figure 7 and Figure 8, Figure 9 further illustrates that nnU-Net 2D produces fragmented segmentation and less accurate boundaries.

**Figure 9.**
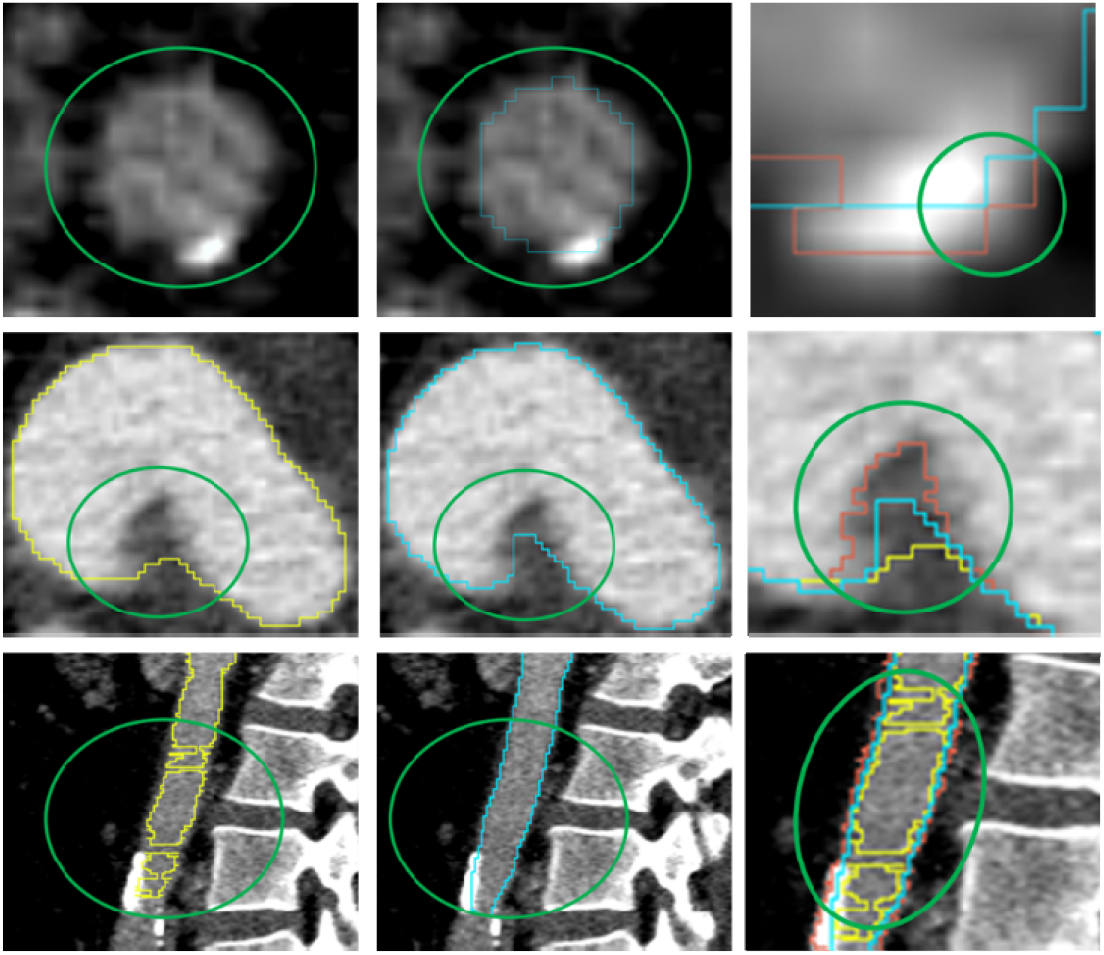
A zoom-in view of the comparison of nnU-Net 2D segmentation (Yellow), nnU-Net 3D segmentation (Blue), and ground-truth (GT) segmentation (Red).

## 4. DISCUSSIONS AND CONCLUSION

In this study, we investigated a deep learning method based on nnU-Net for the segmentation of cardiac CT images to obtain the geometry of the human aorta. We trained both nnU-Net 2D and nnU-Net 3D on the same CT image data. We applied various evaluation metrics to assess the segmentation results from nnU-Nets on the test set. The evaluation results, using DSC, HD95, Jaccard Index, precision, and accuracy, indicate that nnU-Net 3D (i.e., nnU-Net with the 3D U-Net) performs better than nnU-Net 2D (i.e., nnU-Net with the 2D U-Net). However, the recall metric suggests that nnU-Net 2D outperforms nnU-Net 3D. This discrepancy may be caused by the limitations of the dataset size and the imbalanced anatomy of the aorta in our dataset.

As shown in Figures 7, 8, and 9, the segmentation from nnU-Net 2D has larger errors in the bottom section of the aorta. This is mainly caused by the limitation of the 2D U-Net: it generates 3D segmentation results by combining predictions from individual 2D slices, which hampers its ability to capture spatial information between slices. Consequently, segmentation based on single-slice CT images leads to discontinuities in the results. In contrast, the segmentation from nnU-Net 3D demonstrates smoother and more continuous edge contours, reflecting a more accurate representation of the aorta and aortic root compared to nnU-Net 2D.

When constructing the network architecture of nnU-Net 3D, input patches of size 128×128×128 were used, with a voxel spacing of 1.0×0.88×0.88 mm. This configuration ensures that the network’s receptive field can involve the critical features of the aorta and aortic root shapes necessary for effective segmentation.

Although nnU-Net 3D achieves very good results measured by DSC (average DSC > 95%), its HD95 is still large (Max HD95 =18.864mm), indicating that the segmentation of the aortic wall is not accurate enough. Thus, there is still large room for improvement.

The nnU-Net in this study serves as a baseline for developing advanced deep neural networks for more accurate geometry reconstruction of human aorta. Specifically, we will incorporate transformer/attention modules [36,37] into a 3D U-Net to develop novel network architectures. The reconstructed geometries of human aorta can be used for a variety of applications such as stress analysis, hemodynamic analysis, surgical planning, etc., to improve patient outcomes.

## ACKNOWLEDGEMENT

This work was supported in part by the NIH grant R01HL158829.

## Notes

### Competing Interest Statement

The authors have declared no competing interest.

### Summary of Updates

change license to CC-BY so that the preprint may be index in PMC

